# Directional cell motility facilitates side-branching in the mammary epithelium in a tension-sensitive manner

**DOI:** 10.1101/2025.07.30.667704

**Authors:** Beata Kaczyńska, Qiang Lan, Marja L. Mikkola, Satu-Marja Myllymäki

## Abstract

Branching morphogenesis is a fundamental developmental process that shapes the architecture of many epithelial organs, including the mammary gland. In essence, it enables epithelia with cylindrical geometry to grow in a tree-like pattern through repeated cycles of elongation and branching. In the mammary gland, branches form via tip splitting (bifurcation) and budding from the stalk (side-branching). Here, we utilize 3D confocal timelapse imaging of *ex vivo* cultured embryonic mammary glands and a variety of mouse models, to elucidate for the first time, how side branches form in the mammary epithelium. Our results show that branch formation is preceded by outward directional cell migration, whereas cell divisions or their orientation have a more minor contribution. We also demonstrate that frequency of branching increases substantially upon relaxation of epithelial actomyosin network, induced by deletion of *Myh9*. Interestingly, mosaic *Myh9* ablation impedes cell movement amongst neighbors with higher contractility, suggesting that balancing of NMIIA-generated forces are needed for effective collective migration.

## INTRODUCTION

Organs such as lungs, kidneys and exocrine glands, including mammary and salivary glands, have a branched tubular structure that maximizes epithelial surface area for efficient transport or secretory function (Spurlin & Nelson, 2017, Lang et al., 2021). These epithelial surfaces are generated via branching morphogenesis, a process where an epithelial rudiment elongates and branches repeatedly to give rise to a tree-like structure (Myllymäki & Mikkola, 2019).

Epithelial branching is guided by interactions with the surrounding stroma, mediated by paracrine signals and mechanical forces (Lang et al., 2021; Paramore et al., 2022). New branch points can form either through terminal bifurcation (tip splitting), which occurs in most organs, or side-branching (lateral budding from existing ducts), which is detected in lungs, prostate, and mammary glands (Goodwin & Nelson, 2020). Side-branches are quintessential for mammary gland function, as they form during pregnancy and serve as the origin for milk-producing alveoli (Oakes, et al., 2006). However, these branches appear also earlier in development and are thus important for laying down the structural foundation (Satta et al., 2024). Despite their importance, their formation is unclear and may withhold key insights into ductal tumorigenesis (Espina & Liotta, 2011; Ciwinska et al., 2024).

Murine mammary gland development starts at the embryonic ectoderm generating five pairs of mammary buds. Around embryonic day 15 (E15) the buds sprout into the underlying mesenchyme and begin to branch a day later, followed by lumen formation around birth (Myllymäki et al., 2025). Branches form stochastically, but the overall spatial frequency of branching is remarkably even (Satta et al., 2024). Time-lapse imaging has revealed that side-branching accounts for approximately 70% of branching events (Satta et al., 2024). This ability is intrinsic to the mammary epithelium, not specified by the stromal milieu (Lan et al., 2024). After birth, mammary gland grows isometrically with the body until puberty, when reproductive hormones (mainly estradiol) provoke extensive growth and branching, primarily through tip bifurcations (Scheele et al., 2017). Whether additional side-branching occurs also during this stage or is halted until adulthood when induced by progesterone, remains unclear (Brisken & Scabia, 2020).

The molecular cues regulating mammary side-branching are rather well-described (Brisken & Scabia, 2020), but the cellular mechanisms remain unclear. In the single layered epithelium of the developing lung, time-lapse imaging of murine lung explants implicated localized proliferation and oriented cell divisions in side-branch formation (Schnatwinkel & Niswander 2013). In chick lung however, side-branches initiate even if proliferation is blocked, their occurrence depending on actomyosin-driven apical constriction instead (Kim et al., 2013). In the mouse, non-muscle myosin II (NMII) activation was shown to be dispensable (Schnatwinkel & Niswander 2013), although cell shape changes have been observed upon side-branching (Kadzik et al., 2014). Mammary side-branching remains unexplored, but the process of tip bifurcation has been described in more detail (Myllymäki et al., 2023). It involves localized suppression of proliferation and cell motility at the branch point, while cells in the “daughter” tips remain motile/proliferative and change their direction of movement. These cellular dynamics are accompanied by spatially patterned extracellular signal-regulated kinase (ERK) signaling and elevated NMII activity at the leading fronts of tips (Myllymäki et al., 2023).

Here, our aim was to identify the cell behaviors driving side-branching in the mammary epithelium utilizing high resolution time-lapse imaging of ex vivo organ cultures that support vivo-like branch patterning (Satta et al., 2024). We show that side-branches arise in areas of high cellular motility and are preceded by directional cell movement, suggesting that branches initiate much the same way they are elongated (Myllymäki et al., 2023). Branch initiation is sensitive to the levels of actomyosin tension in the tissue, as deletion of *Myh9*, encoding one of three heavy chains of non-muscle myosin II, led to increased branching activity. This applied equally to side-branching and terminal bifurcation, suggesting that they share a common tension-sensitive component. Interestingly, the motility of individual *Myh9*- deleted cells was restricted in a mosaic epithelium, suggesting that collective motility is coordinated by the levels of actomyosin tension.

## RESULTS AND DISCUSSION

### Localized proliferation or oriented cell divisions are unlikely to drive side-branching

The tips of elongating mammary branches are enriched in proliferative cells (Myllymäki et al. 2023) and proliferation marks side-branches in the adult mammary gland (Brisken & Scabia, 2020). To assess the role of proliferation in side-branch formation, we examined cell cycle distribution in a live imaging dataset of ex vivo cultured embryonic mammary glands that express the Fucci2a dual fluorescent cell cycle reporter in a cell cycle phase dependent manner (S/G2/M = hGeminin-mVenus; G0/G1 = hCdt1-mCherry) (Fig. 1A, Supplementary Video 1, Myllymäki et al., 2023). We observed a slight positive trend in cells progressing to the S/G2/M phase of the cell cycle within the side-branching region relative to the non-branching reference region (Fig. 1B, 1C). However, the proportion of S/G2/M cells was not significantly higher, except after the side-branch was already morphologically visible (Fig. 1C) suggesting that a local burst of proliferation is unlikely to drive branch point formation. Unlike post-natal ductal structures, embryonic mammary ducts are relatively proliferative to begin with, where stark differences in cell cycle activity are unexpected. In a fully cellular epithelium of the embryonic mammary gland, changing the geometry of the duct could in principle be accomplished without significantly increasing the local cell number.

**Figure 1.**
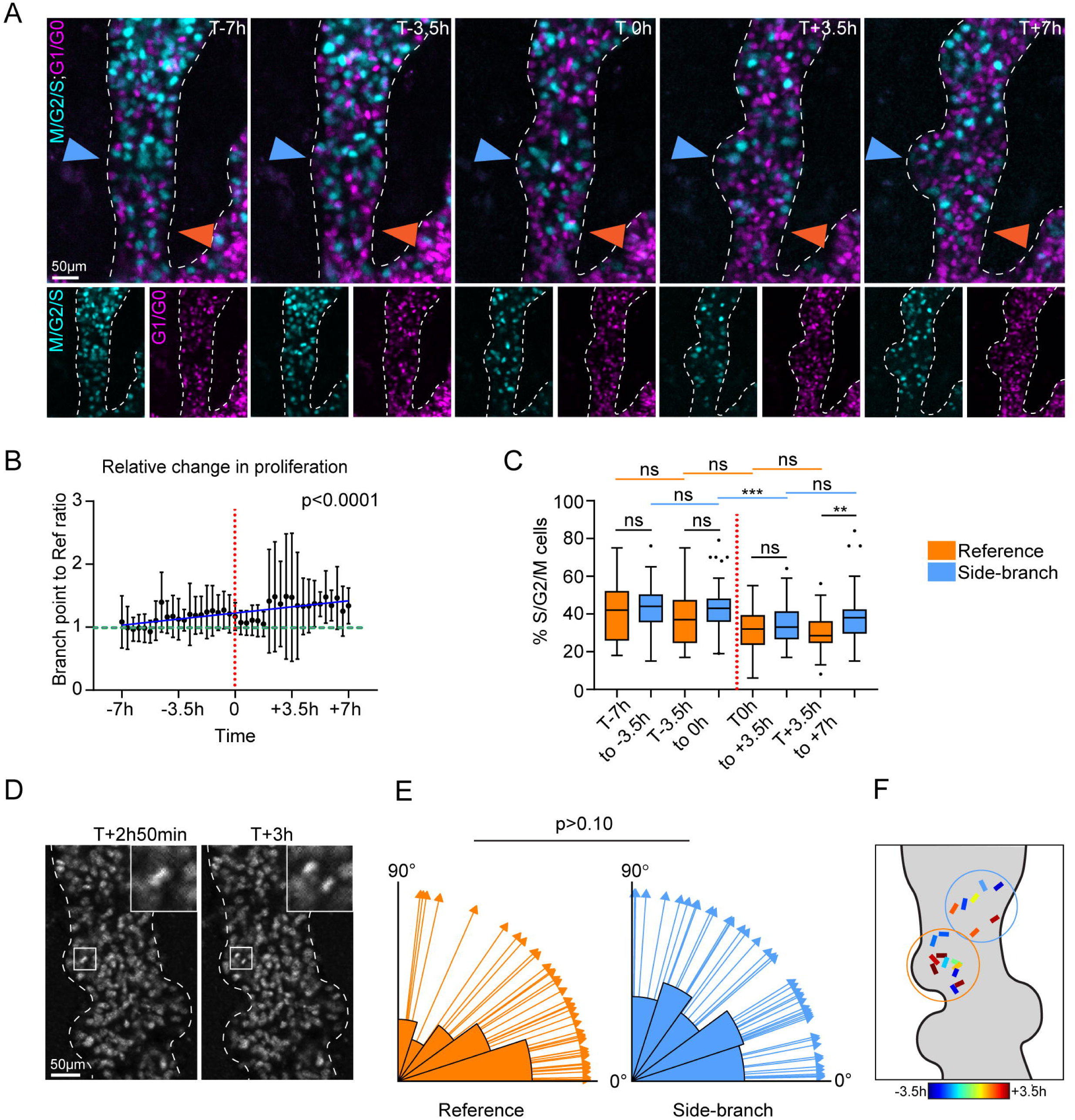
Local changes of proliferation and directional cell divisions are unlikely to drive side-branching **(A)** Optical sections from confocal 4D time-lapse imaging series of mammary glands expressing epithelial Fucci2a cell cycle reporter. Side-branch, blue arrowhead; reference region, orange arrowhead. **(B)** Cell cycle status within the side-branching region relative to the neighboring reference duct, represented as the ratio of % M/G2/S phase nuclei as a function of time, where T0 represents the first emergence of the side branch (n = 8 side-branches from 4 experiments). Correlation between variables was assessed with the Pearson coefficient (R). **(C)** The percentage of M/G2/S cells at binned time windows: -7h to -3.5h (n^ref^=78, n^s-b^=78), -3.5h to 0h (n^ref^=80, n^s-b^=80), 0h to +3.5h (n^ref^= 88, n^s-b^=88), and +3.5h to +7h (n^ref^=80, n^s-b^= 80). Data shown is median with 25th and 75th percentiles (hinges) plus 1.5× interquartile ranges (whiskers). Statistical significance was assessed with Kruskal-Wallis test. **(D)** Orientation of cell divisions was assessed by time-lapse imaging of a constitutive nuclear fluorescent reporter from *K14-Cre+*; *R26R-RG^fl/wt^* mammary glands and measuring the positions of post-metaphase nuclei (right insert). **(E)** Rose diagrams presenting angles of cell divisions in the range of 0-90°, where 90° is perpendicular and 0° parallel to the main duct. Data are pooled from 8 side-branching events, compared to the reference duct (3 experiments; n^ref^=53, n^side-branch^=55). Uniformity of angles was tested with the Rayleigh test (p-value ^ref^ = 5e-04; p-value ^side-branch^= 0.4224) and the difference between the side-branches and reference region with the Watson’s Two-Sample Test of Homogeneity. **(E)** Graphical representation of cell division direction during side-branch formation, side branch area in blue, reference region in orange.

In developing lungs cells change the orientation of division to direct side-branch outgrowth (Schnatwinkel & Niswander, 2013; Kim et al., 2013). In the mammary epithelium, ductal cells divide parallel to the basement membrane whilst cells in stratified ductal tips have been shown to divide more perpendicularly (Huebner et al., 2014). Such a switch in cell division angle might in principle trigger symmetry breaking of the duct. To address this possibility, we utilized the fluorescent R26R-R/G reporter that enables tracking of cells through the cell cycle based on constitutive nuclear mCherry expression (Shioi, et al., 2011) (Fig. 1D). The orientation of cell divisions in the side-branching region did not significantly differ compared to the neighboring, non-branching duct (Fig. 1E). Yet, cells in the duct show stronger preference to divide along the ductal axis, whereas those in the side-branching area seem to divide more randomly (Fig. 1E). Notably, we detected only a few divisions (∼5) per branch prior to branch appearance and their occurrence was non-synchronized (Fig. 1F). Therefore, unless we assume that most escape our detection, these few divisions are unlikely to significantly impact the geometry of the duct. Furthermore, side-branches can be initiated in Mitomycin C-treated embryonic explants (Myllymäki et al., 2023), suggesting that pre-existing cells can accommodate the beginnings of a side branch.

### Directional cell movements contribute to side-branch formation

Based on our previous findings on elongating branches, we hypothesized that cells in the duct may need to become motile and alter the trajectory of their movement towards the direction of side-branch outgrowth. To investigate this possibility, we tracked cell movement in our Fucci2a dataset (Myllymäki et al., 2023), examining side-branching events over a period of 14hrs (Fig. 1A). We found that cells in the side-branching area are faster than those in the neighboring duct well before any morphological sign of a new branch (Fig. 2A, 2B). In contrast, the straightness of the cellś migratory path remained unchanged (Fig. 2C). These results imply that side-branches arise in areas of high motility, likely reflecting the mechanical state of the tissue (Mao & Wickström 2024). Fluid-like behavior in a duct could be enforced by biochemical signals and/or changes in cell density or tension. ERK signaling is an important regulator of collective motion (Lavoie et al., 2020), which we and others have previously observed to be enriched in branch tips, correlating with migratory activity (Huebner et al., 2016, Myllymäki et al., 2023).

**Figure 2.**
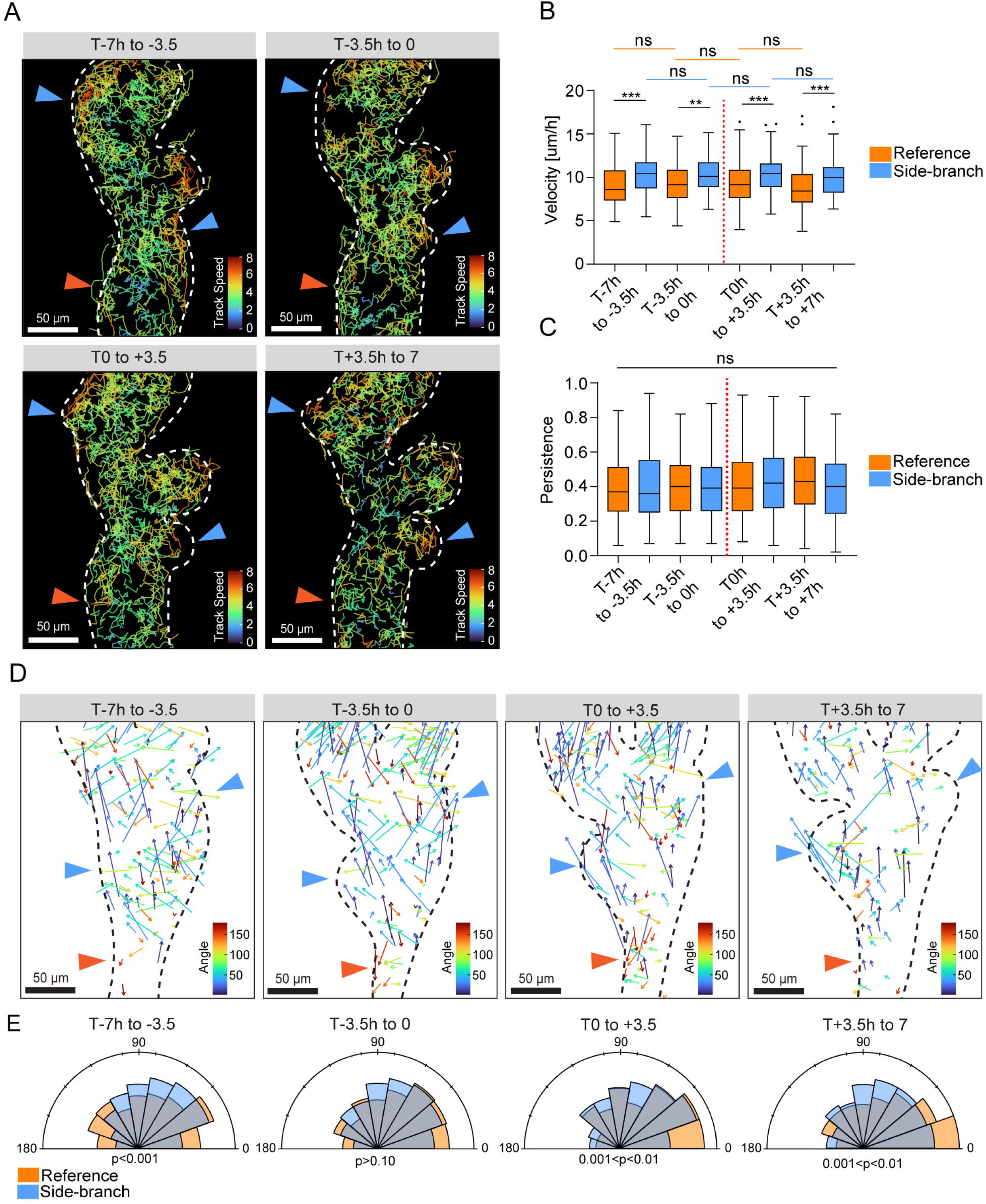
Directional cell movement plays an important role in side-branch formation **(A)** Cell tracks derived from Fucci2a reporter expressing epithelial cells visualized at different stages of side-branching with track velocity color-coded, side-branches (blue arrowheads) and reference region (orange arrowhead). Mean velocity **(B)** and persistence **(C)** were compared between side-branching region and the reference duct (n^ref-7to-3.5^= 163, n^ref-^3.5to0_= 139, n_ref0to+3.5_= 175, n_ref+3.5to+7_= 162, n_s-b-7to-3.5_= 192, n_s-b-3.5to0_= 139, n_s-b0to+3.5_= 158, n_s-^b+3.5to+7^= 114, from 8 side-branching events, 4 experiments). Data shown is median with 25th and 75th percentiles (hinges) plus 1.5× interquartile ranges (whiskers). Statistical significance was assessed with Kruskal-Wallis test. **(D)** Displacement vectors with angles relative to a reference vector running along the duct are shown within side-branches (blue arrowheads) and reference region (orange arrowhead). **(E)** Rose diagrams of cell displacement angles between side-branching and reference region (90⁰ = perpendicular, 0⁰ = parallel/forward, 180⁰ = parallel/backward). Uniformity of angles was tested with the Rayleigh test (p-value ^ref-7to-3.5^= 2e-04; p-value ^ref-3.5to0^= 0; p-value ^ref0to+3.5^= 0; p-value ^ref+3.5to+7^= 0; p-value ^s-b-7to-3.5^= 0; p- value ^s-b-3.5to0^= 0; p-value ^s-b0to+3.5^= 0; p-value ^s-b+3.5to+7^= 2e-04) and the difference between the side-branches and the reference region with the Watson’s Two-Sample Test of Homogeneity.

Next, we explored whether cells also change the direction of movement upon side-branching and by analyzing cell displacement vectors, we found that cells exhibit preferential movement towards the direction of the side-branch 3.5-7h before branching takes place (Fig. 2D-E). Side-branches are limited by the presence of pre-existing branches (Satta et al., 2024), suggesting that repulsive self-avoidance cues may play a role in orienting cell motility.

Indeed, branches avoid beads impregnated by TGF-β ex vivo and in vivo (Silberstein & Daniel 1987, Satta et al., 2024) and TGF-β signaling regulates cell motility in organoids, but its presence at branch points remains unconfirmed (Neumann et al., 2023). In principle, stromal guidance cues could also direct mammary cell motility, but their presence remains unconfirmed as Fgf10 stimulates branching morphogenesis in explants when supplemented into the culture media (Satta et al., 2024, Carabaña et al., 2024), consistent with its uniform expression pattern in vivo (Parsa et al., 2008). Interestingly, mosaic activation of MAPK signaling leads to branching in mammary organoids even in the absence of morphogens (Huebner et al., 2016). Whether such signaling activities coincide with side-branching awaits further studies utilizing fluorescent reporters of ERK activity.

### Loss of non-muscle myosin IIA in mammary epithelial cells leads to impaired movement and decreased directionality

Actomyosin contractility is a fundamental driver of tissue morphogenesis that at the cell scale generates cortical tension, enforces cell junctions, and powers motility (Siedlik & Nelson, 2015). In epithelial tissues, contractility is mediated by the three isoforms of non-muscle myosin II (NMII), NMIIA, NMIIB, and NMIIC. The heavy chain of NMIIA, encoded by the *Myh9* gene, is most abundantly expressed in embryonic epidermis (Ma et al., 2010) that also gives rise to the mammary epithelium. To investigate how *Myh9* deletion impacts cell behavior at a single cell level we performed mosaic, sparse deletion of the *Myh9* gene under Lgr5-EGFP-CreERT2 reporter activated with 4-hydroxytamoxifen (4-OHT) with cytosolic TdTomato as a marker of cells expressing activated CreERT2 and followed the cells for 6 hours by ex vivo imaging (Fig. 3A, Supplementary Video 2 and 3). Analysis of cell tracks revealed that *Myh9*-deleted (TdTomato+) cells were slower (Fig. 3B-C) and moved with less persistence (Fig. 3D) compared to their control, heterozygous counterparts. In addition, *Myh9* cKO cells show loss of directional movement (Fig. 3E-F). These results indicate that at the single cell level, loss of NMIIA-mediated contractility abrogates cellś ability to migrate collectively.

**Figure 3.**
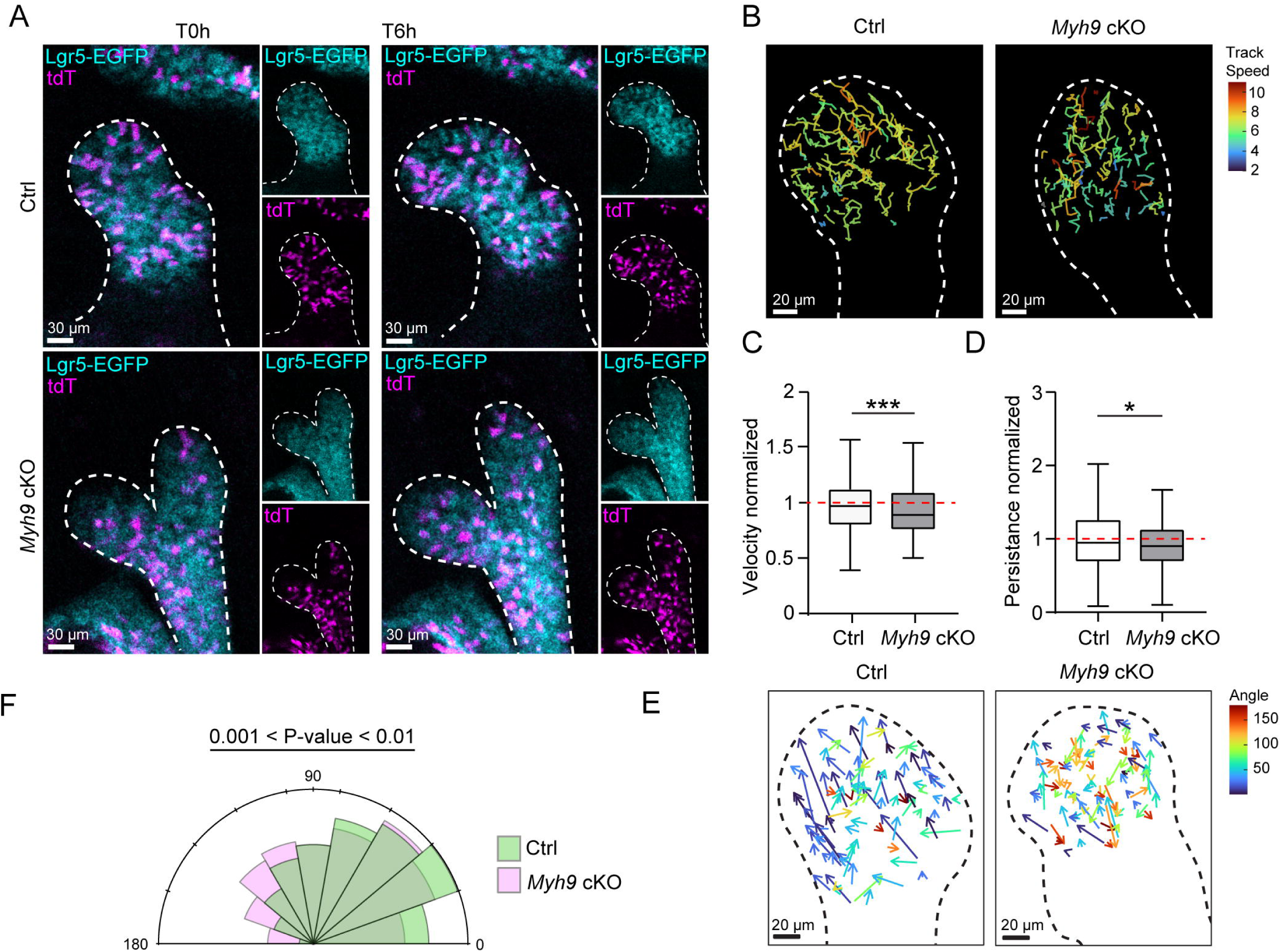
Mosaic loss of NMIIA in mammary epithelial cells impairs motility and directionality **(A)** Confocal slices of 6-hour 4D imaging of 4-OHT treated *Lgr5^EGFP-CreERT2/wt^*; *R26R- TdTomato^fl/wt^* control (*Myh9^fl/wt^*) and sparsely *Myh9* deleted (*Myh9^fl/fl^*) mammary gland. **(B-D)** The velocity **(B, C)** and persistence **(D)** of control (n=498) and *Myh9* deleted (n=327) cells. Data was normalized to the mean of control and represented as median with 25th and 75th percentiles (hinges) plus 1.5× interquartile ranges (whiskers). Statistical significance was assessed with Mann-Whitney test. **(E-F)** Cell displacement vectors of *Myh9^fl/wt^* cells and *Myh9^fl/fl^* cells are shown with their angle relative to the tip displacement color-coded **(E)** and as a rose diagram **(F)** (0⁰ = parallel/forward, 90⁰ = perpendicular, 180⁰ = parallel/backward). Uniformity of angles was tested with the Rayleigh test (p-value^Ctrl^ = 0; p-value*^Myh9^*^cKO^ = 0) and the difference between control and *Myh9* cKO angles with the Watson’s Two-Sample Test of Homogeneity.

### Reduction of NMIIA-mediated epithelial contractility leads to higher branching frequency

As cellular motility correlated with side-branch point formation (Fig. 2) and motility was impaired upon *Myh9* deletion (Fig. 3), we asked how contractility contributes to side-branching. To this end, we generated epithelial conditional *Myh9* knockouts using the *K14- Cre* driver (*Myh9* cKO hereafter) (SFig. 1). Staining of non-muscle myosin IIA heavy chain confirmed successful deletion of *Myh9* in the mammary epithelium (SFig. 1A) associated with reduced levels of NMII activity based on phosphorylated myosin light chain (pMLC) staining (SFig. 1B). At E16.5, when branching begins, confocal whole-mount imaging revealed no difference in the number of tip frequency between *Myh9* cKO and WT glands, although cKO glands were slightly smaller than their WT counterparts (Fig. 4A-C), likely attributed to the role of *Myh9* mammary bud outgrowth and invagination (Trela et al., 2020). Unexpectedly, two days later *Myh9* cKO glands had significantly more branch tips (Fig. 4A, D) without a change in the gland size (Fig. 4E), resulting in a “hyperbranching” phenotype. As branching did not begin precociously (Fig. 4A), NMIIA-mediated epithelial contractility seems to constrain the number of branching events. If NMII is blocked more broadly by treatment with blebbistatin, branching activity ceases completely (Myllymäki et al., 2023). *Myh9* deletion reduced the levels of MLC activity only partially (SFig. 1B), suggesting that other NMII isoforms generate contractility in the mammary epithelium. Indeed, *Myh10* (NMIIB) is expressed in the mammary epithelium at E16.5 while *Myh11* (NMIIC) levels are negligible (SFig. 1C) (Satta et al., 2024). Our results suggest that intermediate levels of contractility produce the highest branching frequency.

**Figure 4.**
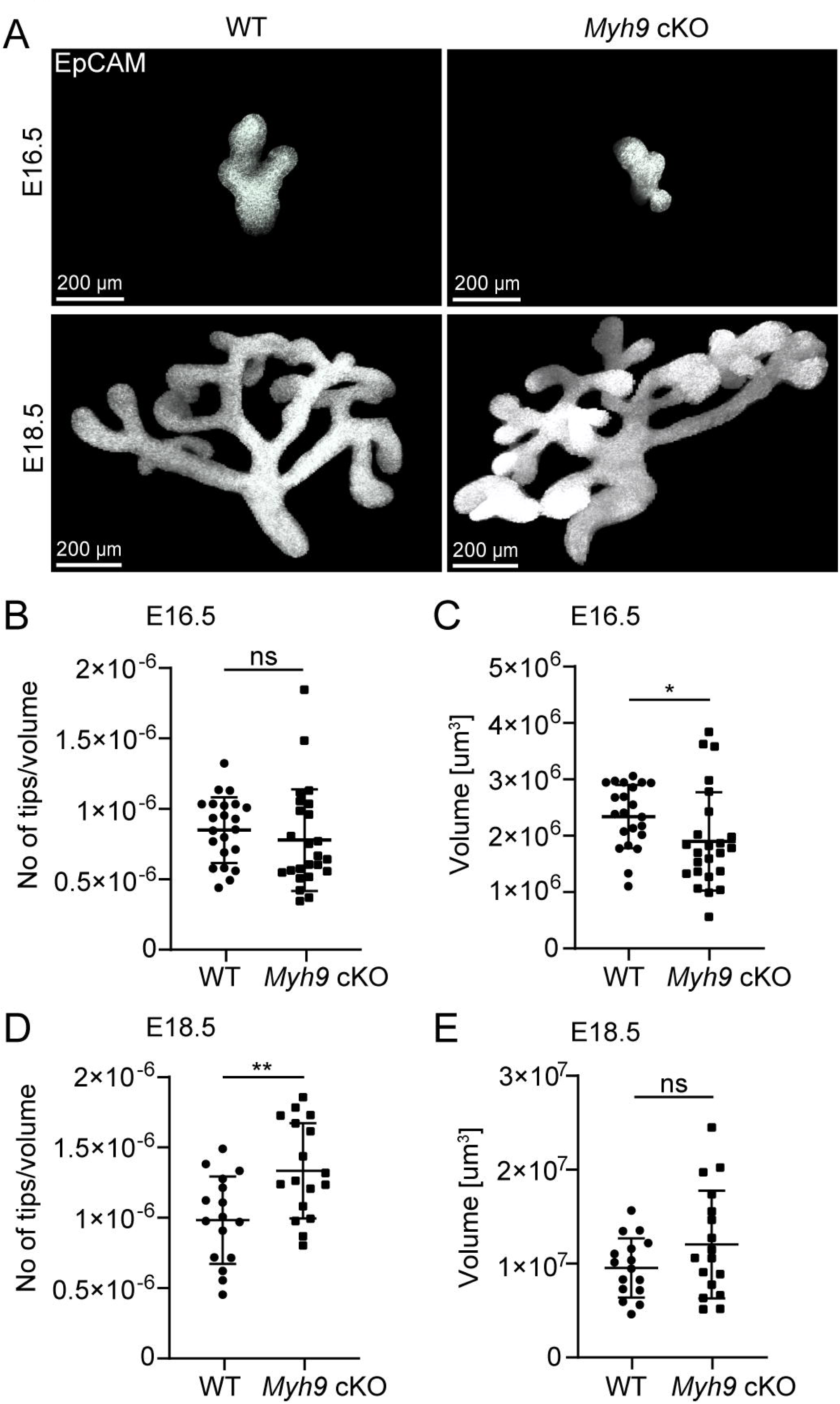
Deletion of *Myh9* in the mammary epithelium enhances branching **(A)** Representative maximum intensity projections of EpCAM-whole-mount stained E16.5 and E18.5 WT and *Myh9* cKO mammary gland (MG) 2. Number of branch tips per volume **(B, D)** and volume **(C, E)** of pooled MG2-3 at E16.5 (n^WT^= 22, n^cKO^= 24) and E18.5 (n^WT^= 16, n^cKO^= 17). Data are shown as mean ± SD; statistical significance was assessed with Mann-Whitney test (B-C; E16.5), or unpaired t-test (D-E; E18.5).

To uncover what type of branching events lead to the observed hyperbranching phenotype, we turned to an ex vivo culture system to follow branching over several days based on a TdTomato Cre reporter (Fig. 5A, Supplementary Video 4 and 5) (Myllymäki et al., 2023; Satta et al., 2024). Our results show that epithelial deletion of *Myh9* gene leads to hyperbranching also ex vivo (Fig. 5A-B), but again, the size of the glands was not significantly affected (Fig. 5C). To our surprise, the proportions of types of branching events did not change (Fig. 5D). It has been postulated before that tip bifurcation and side-branching are fundamentally different forms epithelial remodeling, where tip bifurcation appears to be more reliant upon actomyosin contractility (Wang et al., 2016). Our past and present results imply that the modes of branching have striking similarities in the embryonic mammary gland, possibly because both involve a process of tip “outgrowth” into the stroma, regardless of whether or not a cleft is formed.

**Figure 5.**
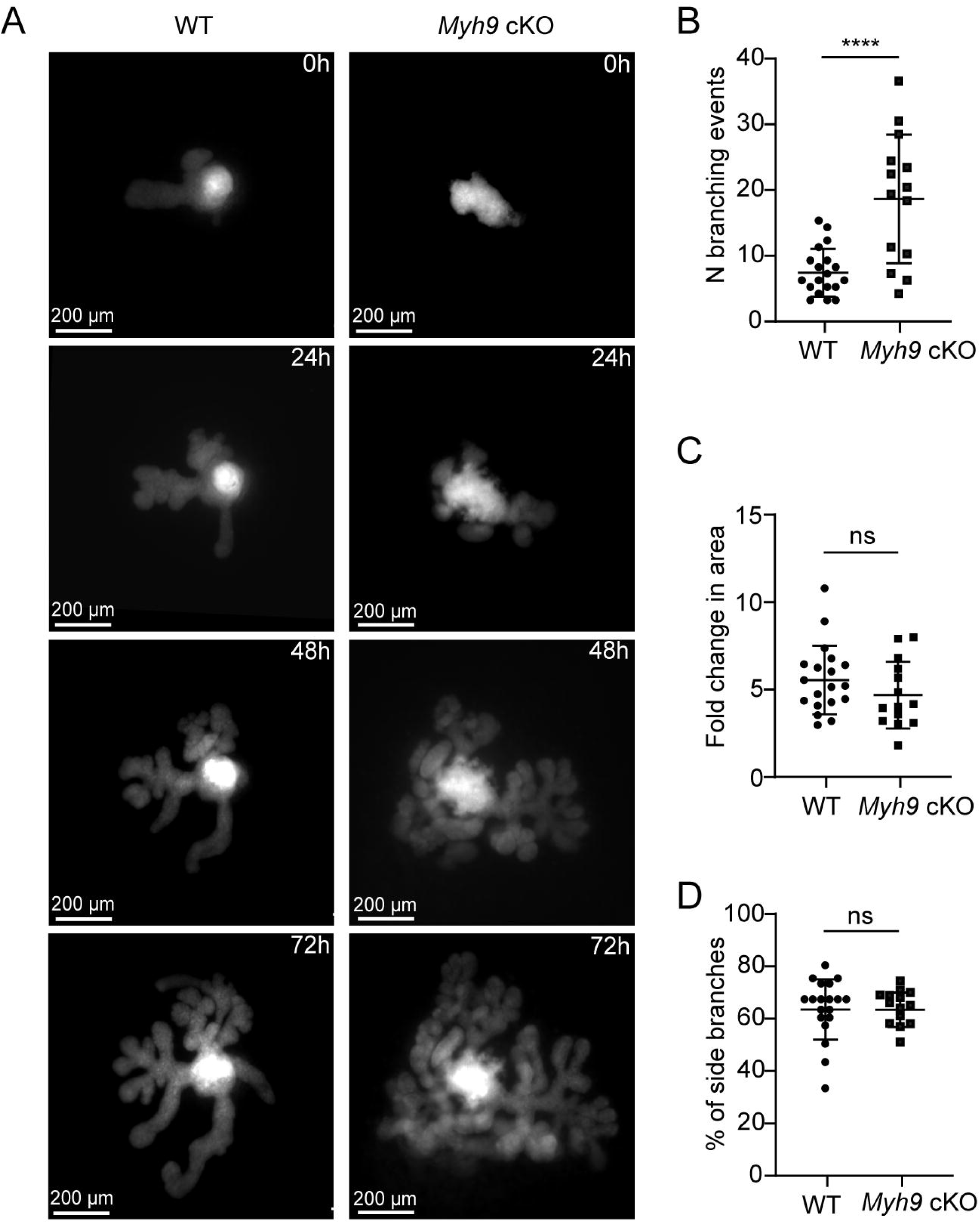
Deletion of *Myh9* in the mammary epithelium does not affect branch mode selection **(A)** A 72-hour time-lapse imaging of ex vivo cultured WT and *Myh9* cKO glands expressing TdTomato Cre reporter. The number of total branching events **(B)**, the fold-increase in gland area **(C)** and percentage of side-branching events **(D)** in WT and *Myh9* cKO ex vivo cultured explants (n^WT^= 19, n^cKO^= 14 glands from 3 experiments). Data are shown as mean ± SD. Statistical significance was assessed with unpaired t-test (B, D), or Mann-Whitney test (C).

Our results on *Myh9* deficient mice demonstrate that global relaxation of actomyosin network, via reduction of NMIIA-mediated contractility, promotes both terminal bifurcation and side-branching events in the mammary epithelium. At the single cell level, *Myh9* deletion reduced cell movement, which seems counterintuitive to the global tissue-level phenotype (Fig. 3A). In the lung, NMII inhibition promotes both branching and migratory behavior in vitro (Plosa et al., 2012). An important distinction in the present study is that the motility of *Myh9* cKO cells was studied in the context of neighbors with higher levels of actomyosin contractility. Collectively migrating epithelial cells remain connected through intercellular junctions and if cortical tension coupled to these junctions is uneven, the “loser” may get left behind. This phenomenon could be related to cell sorting behaviors that are known to be driven by differential cortical tension both in vitro and in vivo (Maître et al., 2012).

Interestingly, NMIIA and NMIIB display functional specificity by regulating different component processes of cell migration and formation of cell-cell junctions (Juanes-Garcia et al., 2015, Heuze et al., 2019). Whether NMII isoforms impart qualitative differences to branching by regulating formation of different types of actomyosin networks in embryonic mammary epithelial cells, awaits further studies on *Myh10* single and *Myh9*/*Myh10* compound mutants.

In summary, our time-lapse imaging data reveals that side-branching in the embryonic mammary gland utilizes directional cell motility. Our analyses on mosaic and global *Myh9* mutants further suggest that collective cell movement requires balancing of NMIIA activity at the cell scale, while high levels of NMIIA restrict branch point formation at the tissue scale.

## MATERIALS AND METHODS

### Ethics statement

All mouse experiments were approved by local ethics committee and the National Animal Experiment Board of Finland (license KEK19-019, KEK22-014). Mice were euthanized with CO2 followed by cervical dislocation.

### Mice

The following mouse strains were maintained in C57BL/6 background: *R26R-Fucci2a-floxed* (Mort et al., 2014), *Lgr5-EGFP-CreERT2* (Barker et al., 2007; The Jackson laboratory, strain 008875), conditional *R26R-RG* (Shioi et al., 2011) and *R26R-TdTomato* mice (The Jackson Laboratory, strain 007914). Transgenic *K14-Cre43* mice (Andl et al., 2004 in NMRI background were used for epithelial expression of Fucci2a conditional fluorescent reporter.

*R26R-RG* mice were crossed with the knock-in *K14-Cre* line (Huelsken et al., 2001) leading to a sparser Cre expression facilitating the tracking of fluorescent cells. *Myh9 floxed* mice (Léon et al., 2007) obtained from the European Mouse Mutant Archive (EM:02572) and *Myh9 floxed*; *R26R-TdTomato-floxed* mice were crossed with the *K14-Cre* mice (Hafner et al., 2004), as previously described (Trela et al., 2021). For mosaic deletion of *Myh9, Myh9^fl/fl^*; *R26R-TdTomato^fl/fl^*females were mated with *Lgr5^EGFP-CreERT2/wt^*;*Myh9^fl/wt^* males. Mice were kept in 12-h light-dark cycles with food and water given ad libitum. The appearance of the vaginal plug was considered as embryonic day (E) 0.5. The embryonic stage was further verified by limb morphogenesis and other external criteria.

### Ex vivo culture of embryonic mammary glands

Embryonic mammary glands were cultured according to an established protocol (Lan et al., 2022). Both flanks of abdominal-thoracic skin containing mammary glands were dissected from E13.5 embryos. The tissues were treated with 1.25 U/ml of Dispase II (4942078001; Sigma-Aldrich) in PBS for 30 min at +4°C and 2 min at room temperature with a pancreatin-trypsin mixture (2.5 mg/ml pancreatin [P3292; Sigma-Aldrich] and 22.5 mg/ml trypsin in Thyrode’s solution, pH 7.4). The enzyme solution was replaced with culture media (10% FBS in 1:1 DMEM/F12 supplemented with 100 µg/ml ascorbic acid, 10 U/ml penicillin and 10 mg/ml streptomycin) and tissues rested on ice for 30 min. The skin epithelium was removed with needles, leaving the mesenchymal tissue with the mammary buds. The tissues were collected on pieces of Nuclepore filter (WHA110605; Whatman) and placed on metal grids in a 3.5 cm Ø plastic Petri dish with culture medium. The explants were cultured up to 7 d in +37°C with air and 5% CO2, replacing the media every other day. 4-hydroxy-tamoxifen (4- OHT, 0.3 µM; H6278, Sigma-Aldrich) was added to the media on day 1 of the culture, left for 1h, and replaced with normal culture media, followed by culture for 5 days prior to confocal imaging.

### Live imaging

#### Wide-field imaging of branching morphogenesis ex vivo

To observe branching morphogenesis by live imaging, mammary glands were dissected from E13.5 *K14-Cre+;Myh9^fl/fl^ ;R26R-tdTomato^fl/wt^* embryos, *K14-Cre+; R26R-tdTomato^fl/wt^*embryos were used as controls. Explants were placed directly into a Trans-well insert to be cultured on a 6-well plate (Falcon). Tissue culture medium was added up to the level of the insert (1.5 ml) and cultured for 4 d in an incubator. Imaging was performed as described previously (Myllymäki et al., 2023). In brief, inverted 3I Marianas microscope equipped with 10×/0.30 N.A. EC Plan-Neofluar Ph1 186 WD = 5.2 M27 at +37°C with 6% CO2. Images were acquired with a LED light source 183 (CoolLED pE2 with 550 nm) and widefield images were acquired with the Andor Neo sCMOS camera operated with Slidebook software at multiple positions at 3-h intervals over a period of 72 h between 4 and 7 d of culture.

Culture medium was replaced every other day.

#### Confocal live imaging

To monitor cell behaviors ex vivo, mammary explants were cultured as previously described (Myllymäki et al., 2023). *K14-Cre;R26R-RG^fl/wt^* or *Lgr5^EGFP-CreERT2/wt^; Myh9^fl/fl^; R26R- TdTomato^fl/wt^* and *Lgr5 ^EGFP-CreERT2/wt^; Myh9^fl/wt^; R26R-TdTomato^fl/wt^* embryos were established as described above and cultured for 4-5d. Thereafter, the explants were transferred on a filter to the bottom of a 5 cm Ø Lumox dish (94.6077.410; Sarstedt) with the tissue side facing down. The edges of the filter were attached to the Lumox dish membrane with Vetbond tissue adhesive (0200742529; 3M Science). After attaching all the explants, 5 ml of culture media was added onto the dish. Imaging was performed with Leica DMI8 inverted laser scanning microscope with an environmental chamber, operated with Las X software. The environmental chamber was set to +37°C air with 5% CO2 and 90% humidity, and the samples were equilibrated there for 1 h before imaging. HC PL APO 20×/0.75 CS2 air objective was used for time-lapse imaging on 4–6 positions at 10 min intervals for up to 24h (for the RG reporter), or for 6 h at 15 min (for 4-OHT treated explants expressing the Lgr5-EGFP-CreERT2 reporter). 3D image stacks were acquired at each position with a 0.5–0.8 µm/px xy resolution and 2 µm z step size, using 600 Hz bi-directional scanning. The pinhole was opened to 2AU to reduce the laser exposure needed. For analysis, videos were cropped and time points limited to include only side-branching events.

### Whole-mount staining and imaging

For whole-mount immunofluorescence staining, dissected ventral skin containing mammary glands were fixed in 4% PFA at 4 °C overnight and washed 4 times 1h in PBS at 4 °C. Samples were blocked with blocking buffer containing 5% normal donkey serum and 0.5% BSA in 0.3% PBST at 4 °C overnight. Primary antibodies were diluted in blocking buffer and incubated with samples for 2–3 d at 4°C. For EpCAM staining, rat anti-mouse CD326 was used as a 1:1,000 dilution (552370; BD Biosciences). After that samples were washed 4 times 1h in PBS at 4 °C and incubated with secondary antibodies and Hoechst (1:2000 dilution, H3570, Thermo Fisher Scientific) diluted in blocking buffer for 2–3 d at 4°C. Secondary antibodies conjugated to Alexa 488 (A21208, Life Technologies), 568 (ab175475, Abcam), and 647 (A48272, Invitrogen) were used in a 1:500 dilution. After washing 4 times 1h in PBS at 4 °C, individual mammary glands were dissected under fluorescence stereomicroscope to remove surplus mesenchymal tissues and mounted on slides using Vectashield (H-1000, Vector Laboratories). The images of whole glands were acquired with Leica Sp8 upright confocal microscope at 1.04 µm intervals with HC PL APO 20×/0.75 IMM CORR CS2 objective with glycerol immersion.

### Immunostaining and imaging of paraffin sections

Samples for sectioning were collected from E18.5 female embryos as described above. Flanks containing mammary glands were fixed in 4% PFA at 4 °C overnight, washed with PBS and processed into paraffin. Prepared samples were then embedded in the paraffin with skin lying flat at the bottom of the mold. Serial sections (5 µm) were cut, deparaffinized and followed by antigen retrieval in TE-buffer (20 mM Tris-HCl, pH 9; 1 mM EDTA) with 0.5% Tween in antigen-unmasking device (Aptum Biologics Ltd). For immunostaining with mouse anti-pMLC (1:300 dilution, 3675S; Cell Signaling Technology), and rat anti-K8 (1:400 dilution, Toma-1-c, DSHB Iowa) antibodies, sections were treated 5 min with 3% H2O2 in PBS, followed by blocking with 10% donkey serum in PBS-0.1%-Tween20 for 1 h at room temperature. To block endogenous mouse IgGs, slides were incubated with the M.O.M blocking reagent (MKB-2213-1, Vector Laboratories) for 1 h at room temperature. After washing three times 5 min with PBS-0.1%-Tween, sections were incubated with primary antibodies overnight at +4°C in PBS-0.1%-Tween20. Incubation with secondary antibodies (1:400 dilution, A21202, Invitrogen; 1:500 dilution, A78946, Life Technologies) and Hoechst (1:2000 dilution, H3570, Thermo Fisher Scientific) was performed at room temperature for 2 h. For immunostaining with Non-muscle myosin heavy chain II-A (1:500 dilution, 909801,

BioLegend), and rat anti-K8 (1:400 dilution, Troma-1-c, DSHB Iowa) antibodies, sections were treated 5 min with 3% H2O2 in PBS, followed by blocking with 10% donkey serum in PBS-0.1%-Tween20 for 1 h at room temperature. After washing three times for 5 min with PBS-0.1%-Tween, sections were incubated with primary antibodies overnight at +4°C in PBS-0.1%-Tween20. Incubation with secondary antibodies (1:500 dilution, A78946, Life Technologies; 1:500 dilution, A31573, Invitrogen) and Hoechst (1:2000 dilution, H3570, Thermo Fisher Scientific) was performed at room temperature for 2 h. The stained samples were imaged with Leica DMI8 inverted laser scanning microscope using 40X/1.25N.A. HC PL APO objective.

### Image analysis

#### Analysis of Fucci2a spots and tracks

Fucci2a live imaging videos (Myllymäki, et al., 2023) were selected based on appearance of side-branching events..The time frame used for analysis included 7h before first morphological changes indicating side-branching event, and 7h after. The area for analysis was selected based on measured length of side-branch that was applied throughout the video. Reference region was chosen in the duct based on lack of any branching events, and the size of the included area was determined based on side-branch length. 6 and 7 µm spots were created in Imaris for G1/G0 and M/G2/S cells, respectively. The % of M/G2/S cells was calculated in each area described above. For cell tracking, the diameter of M/G2/S and G1/G0 nuclei was measured and 8 and 9.3 µm spots were created for G1/G0 and M/G2/S cells, respectively. The position of tracks was determined by their distance to manually placed tissue landmarks (for the side-branch and reference area). The directionality of the cells was analyzed based on their coordinates at the start and end position of the track as displacement vectors. The directionality was determined as the angle between the displacement vector and a reference vector (manually placed in a ductal axis).

#### Cell division tracking in R26R-RG explants

Time lapse videos underwent drift correction in ImageJ utilizing Fast4DReg plugin (Pylvänäinen et al., 2023). Cell divisions were registered manually based on the appearance of the nuclei using multi-point tool. Registered XY positions of the divided nuclei were saved and used to calculate the angle of cell divisions in reference to an axis placed in the main duct.

#### Analysis of fixed whole glands

For mammary gland volume quantification, the border of mammary epithelium was outlined manually based on EpCAM expression, and the surface rendering and volume quantification were performed with Imaris software (version 9.5 or 10.0, Bitplane). The mammary gland tip number was counted manually in 3D. As mammary glands number 2 and 3 were previously found not to significantly differ (Lan et al., 2024), they were pooled for analysis of Myh9 mutants.

#### Quantification of branching events

Live imaging videos in TIFF format were opened in ImageJ and branching events were identified and quantified by visual inspection. Area of the explants were determined by tracing the outline of the epithelium registering the measurement with ROI manager.

#### Analysis of cell behaviors in mice with mosaic Myh9 deletion

Time lapse videos were drift corrected in ImageJ utilizing Fast4DReg plugin. The Lgr5- EGFP-CreERT2 channel was used to create an epithelial surface rendering of a terminal branch in Imaris, spanning 100µm from the leading edge. For tracking of cells inside the branch, the diameter of the TdTomato+ cells was measured and 15 µm spots were created. The position of tracks was determined by measuring the angle of the displacement vectors relative to manually placed reference vector.

#### Data availability

The raw and processed RNA-sequencing data used (Satta et al., 2024) can be found at https://www.ncbi.nlm.nih.gov/geo/query/acc.cgi?acc<u>=</u> GSE236630, hosted at Gene Expression Omnibus.

### Statistics and data representation

All data were analyzed by Prism 9 (GraphPad Software) or R circular package (Agostinelli & Lund, 2022). Statistical tests used are indicated in figure legends. p-values <0.05 were considered significant. Throughout the figure legends: *p<0.05, **p<0.01; ***p<0.001, ****p<0.0001.

## ACKNOWLEDGMENTS

We thank Ms. Raija Savolainen for excellent technical assistance, past and present Mikkola lab members for discussions and advice and Dr. Sara Wickström for support. Imaging was performed at the Light Microscopy Unit of the HiLIFE-Institute of Biotechnology, University of Helsinki. Mouse husbandry was taken care by HiLIFE Laboratory Animal Centre Core Facility, University of Helsinki. This study was financially supported by the Academy of Finland (grant 318287 to M.L. Mikkola), Sigrid Jusélius Foundation (M.L. Mikkola), Finnish Cultural Foundation (B. Kaczyńska), and Ella & Georg Ehrnrooth Foundation (B. Kaczyńska). The funders had no role in study design, data collection and analysis, decision to publish, or preparation of the manuscript.

## AUTHOR CONTRIBUTIONS

Conceptualization: S.-M.M., M.L.M., B.K.; Methodology: B.K., S.-M.M., Q.L.,; Formal analysis: B.K., Q.L., S.-M.M.; Investigation: B.K., Q.L., S.-M.M.; Resources: M.L.M.; Writing - original draft: B.K., S.-M.M., M.L.M.; Visualization: B.K., Q.L., S.-M.M.; Supervision: S.-M.M., M.L.M.; Project administration: S.-M.M., M.L.M.; Funding acquisition: M.L.M.

---

Supplementary Figure 1. Validation of the conditional deletion of *Myh9* in the mammary epithelium **(A, B)** E18.5 WT and *Myh9* cKO mammary glands were stained with Hoechst, keratin 8 (K8), and anti-NMIIA **(A)** or anti-pMLC **(B)**. **(C)** Expression of *Myh9*, *Myh10* and *Myh11* genes measured by RNA-seq in E16.5 mammary epithelium (Satta et al., 2024). Data are shown as mean ± SD.

## REFERENCES

1. Agostinelli, C., Lund, U. (2022). R package ’circular’: Circular Statistics (version 0.4-95). https://r-forge.r-project.org/projects/circular/

2. Andl, T., Ahn, K., Kairo, A., Chu, E. Y., Wine-Lee, L., Reddy, S. T., Croft, N. J., Cebra-Thomas, J. A., Metzger, D., Chambon, P., Lyons, K.M., Mishina, Y., Seykora, J.T., Bryan Crenshaw III, E., Millar, S.E. (2004). Epithelial Bmpr1a regulates differentiation and proliferation in postnatal hair follicles and is essential for tooth development. Development 131, 2257–2268. 10.1242/dev.01125

3. Barker, N., van Es, J.H., Kuipers, J., Kujala, P., van den Born, M., Cozijnsen, M., Haegebarth, A., Korving, J., Begthel, H., Peters, P.J., Clevers H. (2007). Identification of stem cells in small intestine and colon by marker gene Lgr5. Nature 449, 1003–1007. 10.1038/nature06196

4. Brisken, C., Scabia, V. (2020). 90 YEARS OF PROGESTERONE: Progesterone receptor signaling in the normal breast and its implications for cancer. J. Mol. Endocrinol. 65(1):T81–T94. 10.1530/JME-20-0091

5. Carabaña, C., Sun, W., Veludo Ramos, C., Huyghe, M., Perkins, M., Maillot, A., Journot, R., Hartani, F., Faraldo, M. M., Lloyd-Lewis, B., Fre, S. (2024). Spatially distinct epithelial and mesenchymal cell subsets along progressive lineage restriction in the branching embryonic mammary gland. EMBO J. 43: 2308–2336. 10.1038/s44318-024-00115-3

6. Ciwinska, M., Messal, H.A., Hristova, H.R., Lutz, C., Bornes, L., Chalkiadkis, T., Harkes, R., Langedijk, N. S. M., Hutten, S. J., Menezes, R. X., Jonkers, J., Prekovic, S., Grand Challenge PRECISION consortium, Simons, B. D., Scheele, C., van Rheenen, J. (2024). Mechanisms that clear mutations drive field cancerization in mammary tissue. Nature 633, 198–206. 10.1038/s41586-024-07882-3

7. Espina, V., Liotta, L. (2011). What is the malignant nature of human ductal carcinoma in situ? Nat Rev Cancer 11, 68–75. 10.1038/nrc2950

8. Goodwin, K., Nelson, C. M. (2020). Branching morphogenesis. Development 147, dev184499. 10.1242/dev.184499

9. Hafner, M., Wenk, J., Nenci, A., Pasparakis, M., Scharffetter-Kochanek, K., Smyth, N., Peters, T., Kess, D., Holtkötter, O., Shephard, P., Kudlow, J.E., Smola, H., Haase, I., Schippers, A., Krieg, T., Müller W. (2004) Keratin 14 Cre transgenic mice authenticate keratin 14 as an oocyte-expressed protein Genesis 38, 176–181. 10.1002/gene.20016

10. Hartman, J., Mayor, R. (2023). Self-organized collective cell behaviors as design principles for synthetic developmental biology. Semin. Cell Dev. Biol. 141: 63–73. 10.1016/j.semcdb.2022.04.009

11. Heuzé, M. L., Sankara Narayana, G. H. N., DAlessandro, J., Cellerin, V., Dang, T., Williams, D. S., Van Hest, J. C. M., Marcq, P., Mège, R.-M., Ladoux, B. (2019). Myosin II isoforms play distinct roles in adherens junction biogenesis. eLife 8:e46599. 10.7554/eLife.46599

12. Huebner, R. J., Lechler, T., Ewald, A. J. (2014). Developmental stratification of the mammary epithelium occurs through symmetry-breaking vertical divisions of apically positioned luminal cells. Development 141 (5): 1085–1094. 10.1242/dev.103333

13. Huebner, R. J., Neumann, N. M., Ewald, A. J. (2016). Mammary epithelial tubes elongate through MAPK-dependent coordination of cell migration. Development 143 (6): 983–993. 10.1242/dev.127944

14. Huelsken, J., Vogel, R., Erdmann, B., Cotsarelis, G., Birchmeier, W. (2001). β-Catenin Controls Hair Follicle Morphogenesis and Stem Cell Differentiation in the Skin. Cell 105, 533–545. 10.1016/S0092-8674(01)00336-1

15. Juanes-Garcia, A., Chapman, J. R., Aguilar-Cuenca, R., Delgado-Arevalo, C., Hodges, J., Whitmore, L. A., Shabanowitz, J., Hunt, D. F., Horowitz, A. R., Vincente-Manzanares, M. (2015). A regulatory motif in nonmuscle myosin II-B regulates its role in migratory front– back polarity. J. Cell Biol. 209 (1): 23–32. 10.1083/jcb.201407059

16. Kadzik, R. S., Cohen, E. D., Morley, M., Morrisey, E. E. (2014). Wnt ligand/Frizzled 2 receptor signaling regulates tube shape and branch-point formation in the lung through control of epithelial cell shape. Proc. Natl. Acad. Sci. U S A. 111 (34) 12444–12449. 10.1073/pnas.1406639111

17. Kim, H. Y., Varner, V. D., Nelson, C. M. (2013). Apical constriction initiates new bud formation during monopodial branching of the embryonic chicken lung. Development 140, 3146–3155. 10.1242/dev.093682

18. Lan Q., Satta, J., Myllymäki, S.-M., Trela, E., Lindström, R., Kaczyńska, B., Englund, J., Mikkola, M.L. (2022). Protocol for Studying Embryonic Mammary Gland Branching Morphogenesis Ex Vivo. In: Vivanco, M.d. (eds) Mammary Stem Cells. Methods in Molecular Biology, vol 2471. Humana, New York, NY. 10.1007/978-1-0716-2193-6_1

19. Lan, Q., Trela, E., Lindström, R., Satta, J. P., Kaczyńska, B., Christensen, M. M., Holzenberger, M., Jernvall, J., Mikkola, M. L. (2024). Mesenchyme instructs growth while epithelium directs branching in the mouse mammary gland. eLife 13:e93326. 10.7554/eLife.93326

20. Lang, C., Conrad, L., Iber, D. (2021). Organ-Specific Branching Morphogenesis. Front. Cell Dev. Biol. 9:671402. 10.3389/fcell.2021.671402

21. Lavoie, H., Gagnon, J., Therrien, M. (2020). ERK signalling: a master regulator of cell behaviour, life and fate. Nat. Rev. Mol. Cell. Biol. 21(10):607–632. 10.1038/s41580-020-0255-7

22. Léon, C., Eckly, A., Hechler, B., Aleil, B., Freund, M., Ravanat, C., Jourdain, M., Nonne, C., Weber, J., Tiedt, R., Gratacap, M.-P., Severin, S., Cazenave, J.-P., Lanza, F., Skoda, R., Gachet, C. (2007). Megakaryocyte-restricted MYH9 inactivation dramatically affects hemostasis while preserving platelet aggregation and secretion. Blood 110:3183–3191. 10.1182/blood-2007-03-080184

23. Ma, X., Jana, S. S., Conti, M. A., Kawamoto, S., Claycomb, W. C., Adelstein, R. S. (2010). Ablation of nonmuscle myosin II-B and II-C reveals a role for nonmuscle myosin II in cardiac myocyte karyokinesis. Mol. Biol. Cell. 21:3952–3962. 10.1091/mbc.e10-04-0293

24. Maître, J.-L., Berthoumieux, H., Krens, S. F. G., Salbreux, G., Jülicher, F., Paluch, E., Heisenberg, C.-P. (2012). Adhesion Functions in Cell Sorting by Mechanically Coupling the Cortices of Adhering Cells. Science338,253–256. 10.1126/science.1225399

25. Mao, Y., Wickström, S.A. (2024). Mechanical state transitions in the regulation of tissue form and function. Nat Rev Mol Cell Biol 25, 654–670. 10.1038/s41580-024-00719-x

26. Mort, R. L., Ford, M. J., Sakaue-Sawano, A., Lindstrom, N. O., Casadio, A., Douglas, A. T., Keighren, M. A., Hohenstein, P., Miyawaki, A., Jackson, I. J. (2014). Cell Cycle 13 (17), 2681–2696. 10.4161/15384101.2015.945381

27. Myllymäki S.-M., Mikkola M. L. (2019). Inductive signals in branching morphogenesis – lessons from mammary and salivary glands. Curr. Opin. Cell Biol. 61, 72–78. 10.1016/j.ceb.2019.07.001

28. Myllymäki, S.-M., Kaczyńska, B., Lan, Q., Mikkola, M. L. (2023). Spatially coordinated cell cycle activity and motility govern bifurcation of mammary branches. J. Cell. Biol. 222 (9): e202209005. 10.1083/jcb.202209005

29. Myllymäki, S.M., Lan, Q., Mikkola, M.L. (2025) Embryonic Mammary Gland Morphogenesis. Adv Exp Med Biol. 1464:9–27. doi: 10.1007/978-3-031-70875-6_2. PMID: 39821018.

30. Neumann, N. M., Kim, D. M., Huebner, R. J., Ewald, A.J. (2023). Collective cell migration is spatiotemporally regulated during mammary epithelial bifurcation. J. Cell Sci. 136(1): jcs259275. 10.1242/jcs.259275

31. Oakes, S.R., Hilton, H.N., Ormandy, C.J. (2006). Key stages in mammary gland development - The alveolar switch: coordinating the proliferative cues and cell fate decisions that drive the formation of lobuloalveoli from ductal epithelium. Breast Cancer Res 8, 207. 10.1186/bcr1411

32. Paramore, S.V., Goodwin, K., Nelson, C.M. (2022). How to build an epithelial tree. Phys. Biol. 19 061002. 10.1088/1478-3975/ac9e38

33. Parsa, S., Ramasamy, S. K., De Langhe, S., Gupte, V. V., Haigh, J. J., Medina, D., Bellusci, S. (2008). Terminal end bud maintenance in mammary gland is dependent upon FGFR2b signaling. Dev. Biol. 317: 121–131. 10.1016/j.ydbio.2008.02.014

34. Plosa, E. J., Gooding, K. A., Zent, R., Prince, L. S. (2012). Nonmuscle myosin II regulation of lung epithelial morphology. Dev. Dynam. 241:1770–1781, 10.1002/dvdy.23866

35. Pylvänäinen, J.W., Laine, R.F., Saraiva, B.M.S., Ghimire, S., Follain, G., Henriques, R., Jacquemet, G. (2023). Fast4DReg – fast registration of 4D microscopy datasets. J. Cell. Sci. 136 (4): jcs260728. 10.1242/jcs.260728

36. Satta, J. P., Lan, Q., Taketo, M. M., Mikkola, M. L. (2024). Stabilization of Epithelial β- Catenin Compromises Mammary Cell Fate Acquisition and Branching Morphogenesis. J. Investig. Dermatol. 144, 1223e1237. 10.1016/j.jid.2023.11.018

37. Satta, J. P., Lindström, R., Myllymäki, S.-M., Lan, Q., Trela, E., Prunskaite-Hyyryläinen, R., Kaczyńska, B., Voutilainen, M., Kuure, S., Vainio, S. J., Mikkola, M. L. (2024). Exploring the principles of embryonic mammary gland branching morphogenesis. Development 151 (15): dev202179. 10.1242/dev.202179

38. Scheele, C., Hannezo, E., Muraro, M. J., Zomer, A., Langedijk, N., van Oudenaarden, A., Simons, B. D., van Rheenen, J. (2017). Identity and dynamics of mammary stem cells during branching morphogenesis. Nature 542, 313–317. 10.1038/nature21046

39. Schnatwinkel, C., Niswander, L. (2013). Multiparametric image analysis of lung-branching morphogenesis. Dev. Dyn. 242, 622–637. 10.1002/dvdy.23961

40. Shioi, G., Kiyonari, H., Abe, T., Nakao, K., Fujimori, T., Jang, C.-W., Huang, C.-C., Akiyama, H., Behringer, R.R. and Aizawa, S. (2011). A mouse reporter line to conditionally mark nuclei and cell membranes for in vivo live-imaging. Genesis. 49:570–578. 10.1002/dvg.20758

41. Siedlik, M. J., Nelson, C. M. (2015). Regulation of tissue morphodynamics: an important role for actomyosin contractility. Curr. Opin. Cell Biol. 32, 80–85. 10.1016/j.gde.2015.01.002

42. Silberstein, G. B., Daniel, C. W. (1987). Reversible Inhibition of Mammary Gland Growth by Transforming Growth Factor-β. Science 237, 291–293. DOI: 10.1126/science.3474783

43. Sprulin, J. W., Nelson C. M. (2017). Building branched tissue structures: from single cell guidance to coordinated construction. Phil. Trans. R. Soc. B 372:20150527. 10.1098/rstb.2015.0527

44. Trela, E., Lan, Q., Myllymäki, S.-M., Villeneuve, C., Lindström, R., Kumar, V., Wickström, S. A., Mikkola, M. L. (2021). Cell influx and contractile actomyosin force drive mammary bud growth and invagination. J Cell Biol. 220(8):e202008062. 10.1083/jcb.202008062

45. Wang, S., Sekiguchi, R., Daley, W.P., Yamada, K.M. (2017). Patterned cell and matrix dynamics in branching morphogenesis. J. Cell Biol. 216:559–570. 10.1083/jcb.201610048

