## Supplementary figures and images for "Directional cell motility facilitates side-branching in the mammary epithelium in a tension-sensitive manner"

### Figure S1

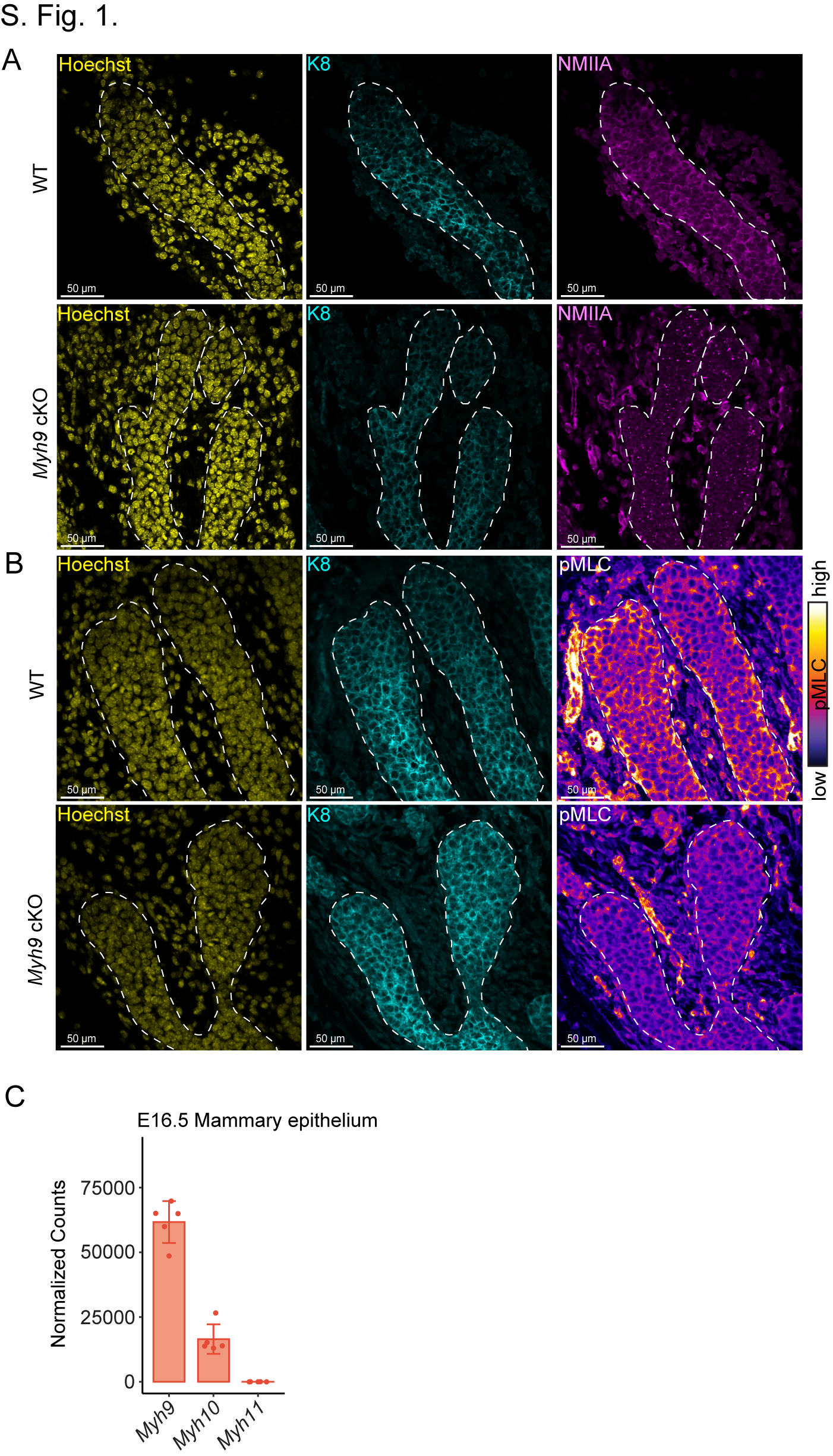
